# Genomic instability in patients with sex determination defects and germ cell cancer

**DOI:** 10.1101/2022.06.08.495249

**Authors:** Maria Krivega, Jutta Zimmer, Anna Slezko, Petra Frank-Herrmann, Julia Rehnitz, Markus Hohenfellner, Markus Bettendorf, Marcin Luzarowski, Thomas Strowitzki

**Affiliations:** Research Group of Gonadal Differentiation and Embryonic Development, Department of Gynecology Endocrinology & Infertility Disorders, Women Hospital, University of Heidelberg, 69120 Heidelberg, Germany; Department of Gynecology, Endocrinology & Infertility Disorders, Women Hospital, University of Heidelberg, 69120 Heidelberg, Germany; Department of Urology, University Hospital Heidelberg, 69120 Heidelberg, Germany; Division of Pediatric Endocrinology, Children’s Hospital, University of Heidelberg, 69120 Heidelberg, Germany; Core Facility for Mass Spectrometry & Proteomics, ZMBH, University of Heidelberg, 69120 Heidelberg, Germany

## Abstract

The ability to transmit genetic information through generations depends on preservation of genome integrity. Genetic abnormalities affect cell differentiation, causing tissue specification defects and cancer. We addressed genomic instability in individuals with Differences of Sex Development (DSD), characterized by gonadal dysgenesis, infertility, high susceptibility for different types of cancer, especially Germ Cell Tumors (GCT), and in men with testicular GCTs. Whole proteome analysis of leukocytes, supported by specific gene expression assessment, and dysgenic gonads characterization, uncovered DNA damage phenotypes with altered innate immune response and autophagy. Further examination of DNA damage response revealed a reliance on deltaTP53, which was compromised by mutations in the transactivation domain in DSD-patients with GCT. Accordingly, drug-induced rescue of DNA damage was achieved by autophagy inhibition but not by stabilization of TP53 in DSD-patients’ blood *in vitro*. This study elucidates possibilities for prophylactic treatments of DSD patients as well as new diagnostic approaches of GCT.

**Teaser:** DNA damage phenotypes accompany aneuploidy of sex chromosomes and link to infertility and high propensity to germ cell tumor development.

## Introduction

Mammalian gonads’ specification is a unique process resulting in two very different organs systems from one bipotential progenitor. It is tightly regulated by a complex gene network providing specific conditions for cell lineages differentiation, followed by gonadal sex determination. Many of these genes are encoded on X and Y chromosomes and expressed in a dose dependent manner(*1*).

It is now widely accepted that Y chromosome is susceptible to DNA deletions which causes spermatogenic failure (*2*). Instability of Y chromosome can be connected to high frequency of repetitive DNA elements, which emphasizes the importance of DNA repair mechanisms to retain integrity of Y chromosome (*3*). However, inability to retrieve lost information because of the non-existent paired chromosome compromises a possibility of highly precise homologous recombination mechanism of DNA repair. As it was recently shown, artificially designed missegregation of Y-chromosome leads to its rapid loss from the cells (*4*). It was accompanied by Y-chromosome isolation and shattering in the micronuclei, followed by an activation of the error prone non-homologous end joining (NHEJ) DNA repair mechanism. NHEJ further provokes genomic instability and increases levels of double stranded brakes (DSB) in DNA, illustrated by highly sensitive epigenetic histone modifications (*5*). The first steps of DNA repair are supported via phosphorylation of Histone protein H2AX on Ser139 (γH2A) to mark DSBs, and of TP53 to promote check points for DNA repair (*6*). This proper mechanism of DNA damage response (DDR) is critical for genomic integrity of the entire cell and can be associated with the stability of the Y chromosome.

Abnormalities related to dysfunctional sex chromosomes are associated to Differences of Sex Development (DSD). In fact, two discoveries regarding DSD had shaped our current understanding of sex determination in humans. Identification of Turner (45,X0, females) and Klinefelter (47,XXY, males) syndromes defined the critical role of Y chromosome for a male differentiation (*7, 8*). And much later, Y chromosome specific SRY-gene translocation was shown to be sufficient to transform XX-individuals into males (*9*). These days DSD is characterized by a large spectrum of complex conditions that among others include dysgenetic gonads (DG), sex reversal, and inability to produce mature germ cells. Besides these severe infertility phenotypes, these individuals have a high risk to develop germ cell tumors (GCT) (*10*).

DGs are immature and contain remnants of primordial germ cells that are unable to differentiate and might lead to gonadoblastoma (GB), a primarily benign tumor with malignant capacity. DSD is described in patients with Swyer syndrome (46,XY, females) that is often accompanied by GBs (*11, 12*). Patients carrying genetic mutations on Y-chromosomes have a 50% risk of developing Germ Cell Neoplasia *in situ* (GCNIS) and GB, and 50% can even develop into malignant tumors (*13*). Alternative cases of 46,XY DSD-patients possess 12-40% cancer risk (*14*). In complete or partial androgen insensitivity syndrome (CAIS, PAIS, 46,XY, females) a malignant development is also increased with a 15% risk for adult women (*15*). Women with Turner Syndrome carrying Y-chromosome material have 12-40% of GNIS/GBs and 3% dysgerminoma, and a risk for ovarian cancer (*16, 17*). Interestingly, Klinefelter patients are known to suffer from mediastinal GCTs (*18*). Strong resistance of these extragonadal tumors in comparison to gonadal GCTs is supported by P53 (*19*). In addition to these observations, non-malignant GB was previously described as P53 positive (*20*). Despite of significant efforts to identify the critical cancerogenic regions on Y-chromosome, the reasons behind the discrepancies among different groups of patients with DSD and the driving force behind GBs transformation remain unclear.

A malignant differentiation of GB in dysgenic gonads was associated with the presence of Y-chromosome material, which might be the easiest target for genotoxic stress, as mentioned above. However, embryonal carcinoma can occur in women with karyotypes 45,XO and 46,XX due to the specific autosomal mutations, without presence of Y-chromosome (*21*–*24*). One should take into consideration, that non-DSD genetic syndromes e.g. Down syndrome (DS, Trisomy 21 chromosome), are often accompanied by delayed germ cell maturation resolving in testicular germ cell tumor (TGCT) (*25*). These observations question malignant activity of unstable Y-chromosome, and rather bring into focus the integrity of the entire genome.

The cellular defects associated with unbalanced karyotypes, including sex chromosome aneuploidy, may be caused by the disruption of cellular homeostasis due to the deregulation of genome-wide expression. Constitutive alterations of these genes cause a whole pleiad of cellular stress phenotypes including, but not limited to proliferation delay, defects in proteostasis, DNA damage and activation of innate immune response (*26*–*29*). For example, Turner syndrome was associated with specific changes in a wide range of genes involved in cellular metabolism, immune response, genome methylation status, embryonic development and morphogenesis (*30*). This leads to susceptibility of DSD-patients to many other non-GCT types of cancer that go beyond reproductive tissues e.g. solid tumors of skin and central neural system (*31*–*33*). These cellular stress phenotypes were previously described in a context of genomic instability and aneuploidy in general (*27, 34, 35*). Therefore, in this work we studied whether different patients’ groups with variations of their gonadal differentiation share common mechanisms which might cause an increased susceptibility to cancer.

## Results

### Increased DNA damage levels in DSD-individuals

Compromised DNA integrity is a well-known sign of genomic instability, the leading cause of cancer. To evaluate general levels of DNA damage we analyzed leukocytes from peripheral blood of 39 different DSD-patients with Swyer, CAIS, Turner and Klinefelter syndromes (Table S1). For control we used blood from fertile men and women. Patients with Swyer syndrome or CAIS patients who underwent gonadectomy for routine histological analysis formed an additional group with histologically proven GCT (DSD-GCT). Since the other two patients’ groups don’t usually undergo gonadectomy due to the established low risk of GCT occurrence, no gonadal tissue from these patients was available for analysis.

DNA damage levels were estimated by assessing an increase of DSB via γH2AX. We identified more than two times upregulated γH2AX relative levels in leukocytes of patients with Swyer syndrome (Fig. 1a-c), similar to those in DSD-patients with GCT. CAIS-samples were not significantly different from a control (Fig. 1c, d). Interestingly, patients with Turner and Klinefelter syndromes showed a pronounced increase by four times (Fig. 1c, e, f). As alternative confirmation of DNA damage we analyzed *H2AX* RNA expression(*36*). We observed transcriptional upregulation which was accompanied by over two folds increase in total H2AX protein levels exclusively in the DSD-GCT group (Fig. 1a,b, d-h).

**Figure 1.**
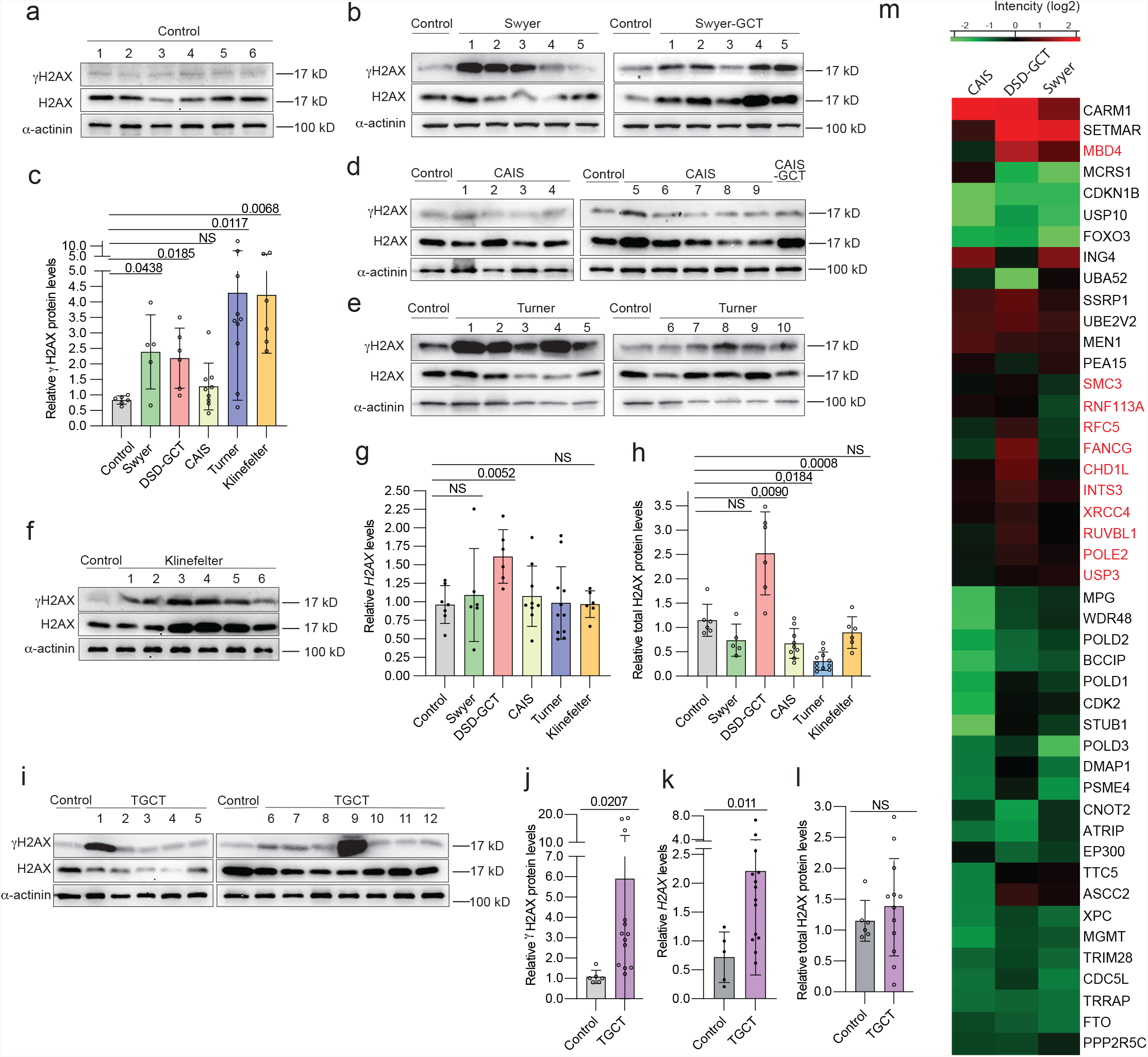
DNA damage in leukocytes of DSD- and TGCT-patients. Immunoblotting for γH2AX and total H2AX proteins from leukocytes of controls (a) and patients with Swyer and Swyer-GCT (b). Quantification of relative γH2AX protein levels calculated for DSD-group (c). Immunoblotting for CAIS and CAIS-GCT (d), Turner (e), Klinefelter (f) syndromes. Relative gene expression of *H2AX* measured for DSD-group (g). Quantification of relative total H2AX protein levels normalized to α-actinin of DSD (h). Immunoblotting for TGCT-group (i) and its relative quantification (j). qRT-PCR for *H2AX* gene expression for TGCT-samples (k). Total H2AX protein levels normalized to α-actinin in TGCT-samples (l). (m) Heat map of GO selection on proteins involved in DNA damage repair from entire proteome of DSD-patients, normalized to control. Unpaired t test was used for statistical analysis, p values are indicated.

To strengthen phenotypes described for DSD-GCT-patients, we included in our study an additional test group of 14 men diagnosed with TGCT (Table S2). The leukocytes from peripheral blood uncovered a strong increase by six times in γH2AX and by two times in *H2AX* transcripts (Fig. 1i-k). In turn a slight increase in total H2AX protein levels was not statistically significant (Fig. 1i,l). Generally, we have not observed any significant correlation in DNA damage markers with increased age of control and patient groups on protein levels (Fig. S1a, b). There was a correlation in control samples for *H2AX*, however the fold changes were not higher than in patient groups (Fig. S1c).

For further evaluation of genomic instability, we performed mass spectrometry analysis for the entire proteome isolated from leukocytes of Swyer, CAIS and DSD-GCT groups. All the patient samples were normalized to control and mainly showed upregulation of immune response and extracellular signaling, however pathways involved in RNA biogenesis were downregulated (Fig. S1d, e; Dataset S1). While there were generally more deregulated processes in GCT presence, we observed a consistent decrease in DNA damage repair proteins in all patients’ groups (Fig. S1e, f). In turn some genes e.g. SMC3, FANCG, CHD1L, XRCC4 were upregulated in DSD-GCT, likely representing requirement of specific DNA repair mechanisms for this patients’ group (Fig. 1m, Dataset S2).

### Dysgenic gonads contain cells with nuclear deterioration

Nuclear deformation and micronucleation illustrate severe problems with genome organization, chromosomal missegregation, aneuploidy and associated DNA damage(*37*). As recently shown germ cell aplasia is also associated with nuclear indentation in gonads of infertile men(*38*). Therefore, we analyzed the shape of the nuclear envelop in biopsied gonadal tissue of five Swyer and six CAIS patients. For control we studied testicular biopsies from fertile men with mature spermatogenesis (Fig. 2a). In contrast to control, patients with Swyer syndrome had misshaped nuclei and multinucleation characterized by decreased circularity and roundness parameters, also with GCT (Fig. 2b-e). AMH-positive Sertoli cells of CAIS-patients looked similar to the control, which reflected on little decrease in nuclei circularity and roundness, likely illustrating absence or little deterioration of genomic integrity (Fig. 2d-g). We also observed increased percentage of γH2AX positive cells, with upregulated number of puncta, in gonads of Swyer, DSD-GCT and TGCT-patients, while CAIS syndrome was associated with little or no change (Fig. 2h-k).

The signs of compromised genome integrity in gonadal tissue supplement our data on peripheral blood and imply presence of systemic problems causing DNA damage in DSD-patients.

**Figure 2.**
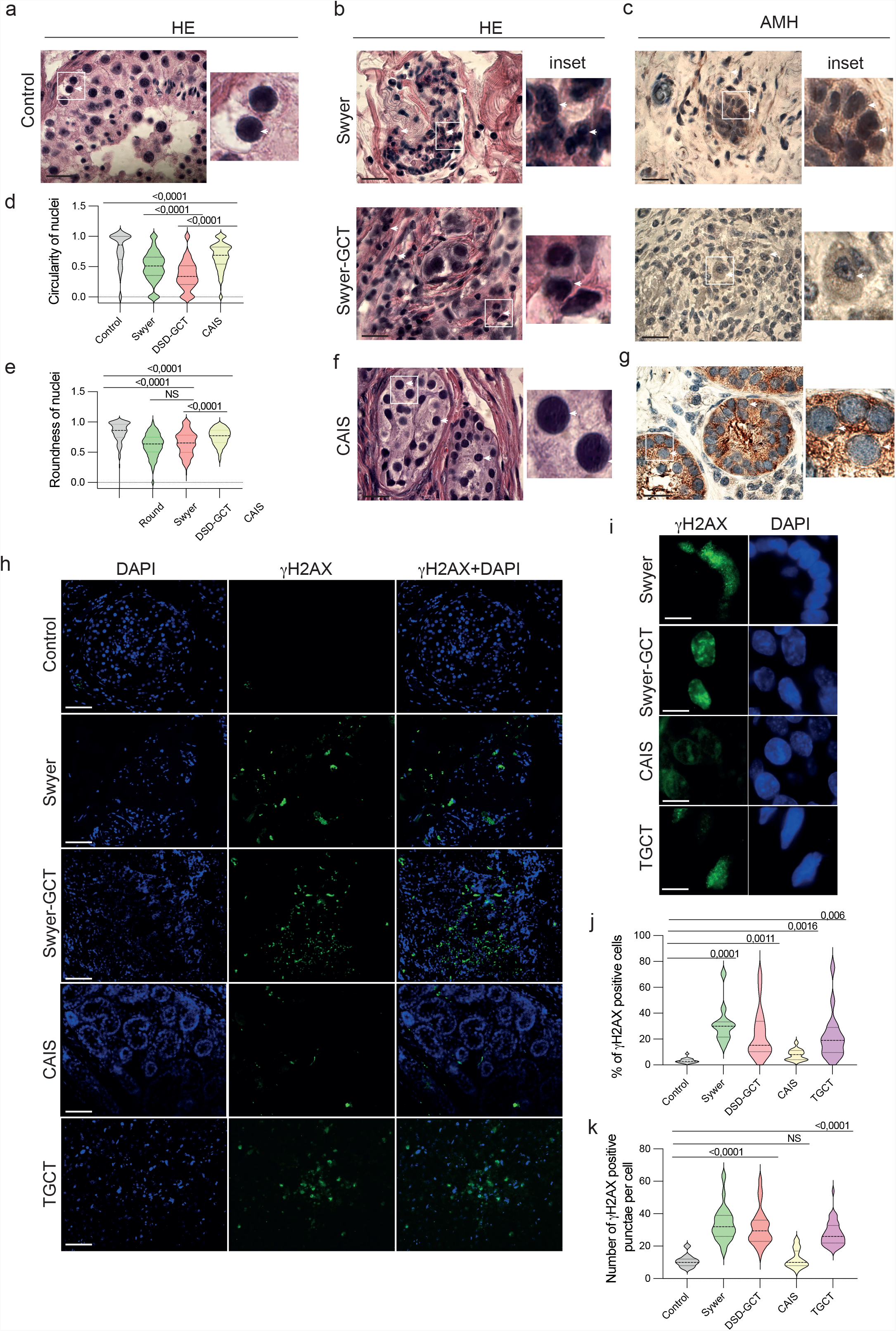
Genotoxic stress phenotype in cells of gonadal tissue from DSD-patients. Gonadal tissue was stained with Hematoxylin-eosin (HE) staining to define nuclei in control (n=210) (a), Swyer (n=342) and Swyer-GCT (n=186) (b). Immunohistochemistry for AMH to define Sertoli cells in Swyer and Swyer-GCT(c). Plots for circularity (d) and roundness (e) of the nuclei in Sertoli cells of DSD-patients. CAIS group (n=526) staining for HE (f) and AMH (g); n-number of cells analyzed per each group. (h, i) Immunofluorescence for γH2AX, positive puncta are indicated by while arrows. (j) Quantification of percentages of γH2AX positive cells in control (n=16), Swyer (n=11), DSD-GCT (n=25), CAIS (n=14), TGCT (n=16); n-number of images analyzed for each patients group. (k) Quantification of number of γH2AX positive cells in control (n=11), Swyer (n=75), DSD-GCT (n=46), CAIS (n=33), TGCT (n=40); n-total number of cells analyzed in each group. Paired t test was used for statistical analysis, p values are indicated. Scale 10 µm (a-c, f, g), 2 µm (h), 0,4 µm (i). n-number of analyzed cells.

### Changes in innate immune response are associated with genotoxic stress in DSD-patients

To estimate whether increased DNA damage has biological significance we examined changes in well-known DDR phenotypes. A number of cytoplasmic responses can be activated to alleviate deleterious consequences of genotoxic stress and support cell survival. Autophagy, regulating protein degradation, and type I Interferon signals were shown to reflect compromised integrity of genomic DNA in multiple cell lines and animal models (*27, 39*). However, nobody has assessed this interplay in patients’ samples.

We showed that wide range of proteins involved in immune response is generally upregulated in DSD-patients’ samples (Fig. S1d, e, S2a). We observed particularly high levels of DDR-relevant immune response proteins in DSD-GCT group, e.g. STING1, IFITM2, IFI44, MX1, MX2 (Fig. 3a, b; Dataset S3). Interestingly RNA helicases like DHX36 and DDX60 were increased as well, potentially implying involvement of RNA regulation in innate immune response induction. Additional proof of DDR in blood of DSD-GCT and TGCT patients was a significant transcriptional upregulation of type I interferon *IFNβ* and interferon stimulating genes (ISG) *ISG15* and *ISG56* up to 15 folds (Fig. 3 c-f). ISGs’ induction was exclusively associated with GCTs, since CAIS-patients with inhibited androgen signaling only upregulated *IFNβ*. Indeed, detailed proteome analysis showed different character of immune response activation between DSD-GCT and CAIS groups (Fig. 3g, h; Dataset S4).

**Figure 3.**
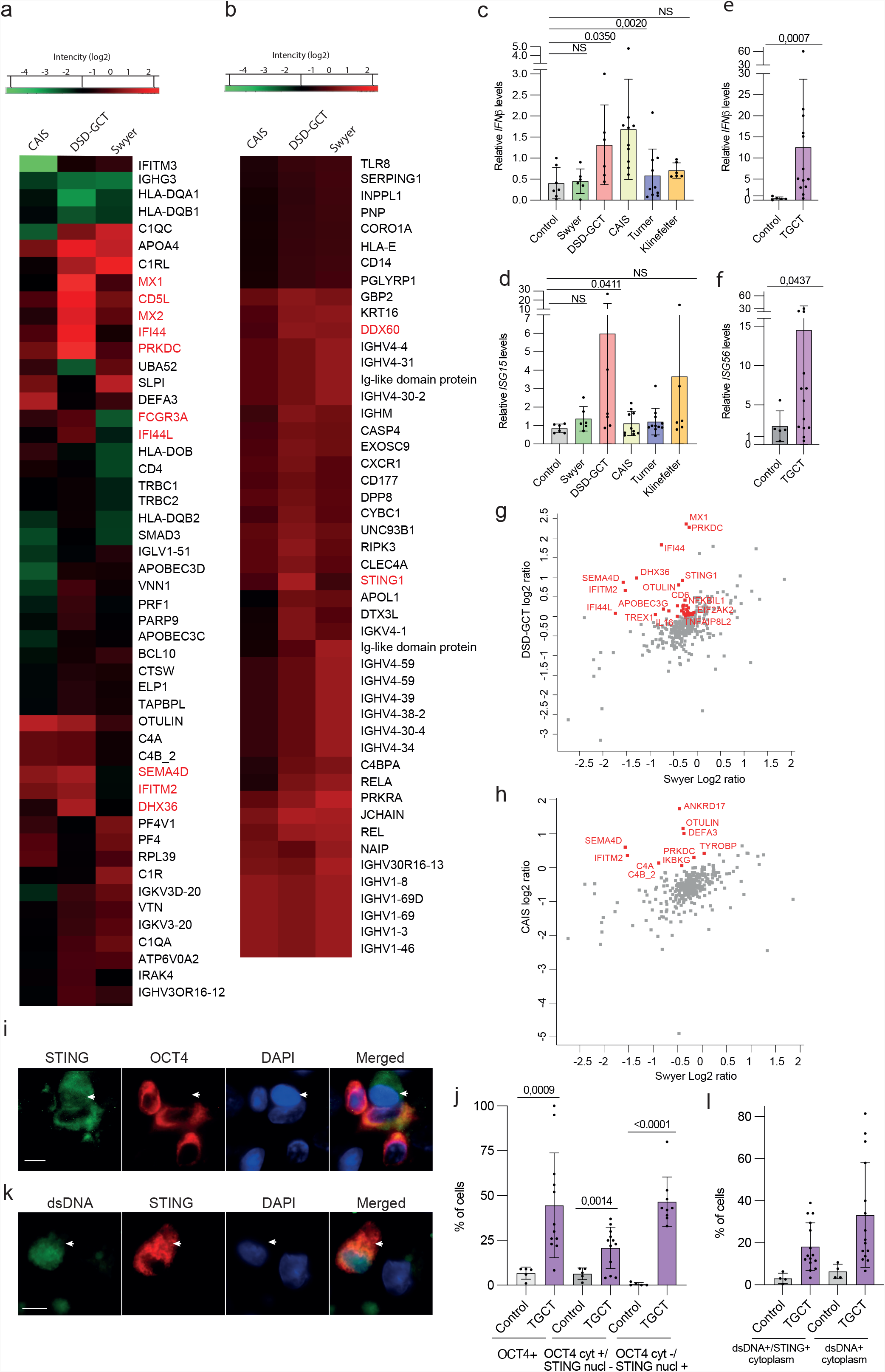
Immune response in DSD and TGCT samples. Heat maps (a, b) and 2D plots (g, h) representing a GO selection of immune response proteins from the entire proteome mass spectrometry data of patients’ leukocytes. qRT-PCR data of innate immune specific gene expression in leukocytes of *IFNβ* (c), *ISG15* (d) for DSD group and *IFNβ* (e), *ISG56* (f) for TGCT group. Immune fluorescence analysis of TGCT gonadal tissue biopsy for STING and OCT4 proteins (I; n(TGCT)=523; n(control)=871; arrow points to OCT4-negative non-GCT cell that has STING in the nucleus, in contrast to other OCT4-positive cells with STING exclusively conzentrated in the cytoplasm) or dsDNA (k; n(TGCT)=2140; n(control)=783; arrow points to colocalization of dsDNA and STING in the cytoplasm) and the quantification (j, l). Unpaired t test was used for statistical analysis, p values are indicated. Scale 0,4 µm (i, k). n-number of analyzed cells. Cyt-cytoplasm; nucl-nucleus.

In turn, *IL6* stimulated by alternative innate immune response pathway of nuclear factor kappa B (NFkB), was not altered in DSD-GCT (Fig. S2b). Importantly, we observed clear positive correlation of *H2AX* and *IFNβ* transcripts in Swyer, Turner and TGCT-individuals (Fig. S2c, d). CAIS with low DNA damage levels didn’t show similar correlation.

STING is critical for stimulating type I IFN response in a context of genomic instability. Therefore, we aimed to connect our finding in peripheral blood leukocytes with GCT cells. We analyzed gonadal tissue samples of TGCT-patients and showed increased presence of OCT4+ GCT cells with active STING protein concentrating in the cytoplasm, where it functions to transmit the signal for innate immunity activation (Fig. 3i, j). While the cells with exclusively nuclear STING were negative for OCT4. Genomic dsDNA leakage from the nucleus is a sign of compromised genome stability. Using the same TGCT tissue samples we observed increased number of cells positive for dsDNA, concentrated together with STING in the cytoplasm (Fig. 3k, l). Taken together, we show upregulated immune response in leukocytes of DSD and TGCT-patients with specific DDR relevant stimulation in patients’ gonads with GCT.

### Autophagy is inhibited in leukocytes of DSD-patients

Autophagy is known to be deregulated upon genomic instability. We showed accumulation of autophagy specific proteome in leukocytes of DSD-patients (Fig. 4a, Fig. S2e; Dataset S5). Generally, CAIS-patients show less accumulation and even some decrease of autophagy-specific proteome, which is consistent with absent DNA damage phenotype in these samples. To understand whether it is caused by inhibited protein degradation or stimulated protein production, we analyzed transcription of autophagy markers *LC3* and *P62* in patients’ leukocytes, and observed statistically significant decrease by two folds in *LC3* transcripts DSD-GCT and TGCT-samples (Fig. 4b-e). Later stages of autophagy require fusion with lysosome for proteolysis, and we didn’t detect significant downregulation in lysosomal-specific gene expression in both GCT-groups, but rather *CTSB* increase in DSD-GCTs when compared to control or Swyer and CAIS-patients (Fig. S2 f-i). These data indicate autophagy regulation on early stages already on transcriptional level in samples with DSD and GCT.

**Figure 4.**
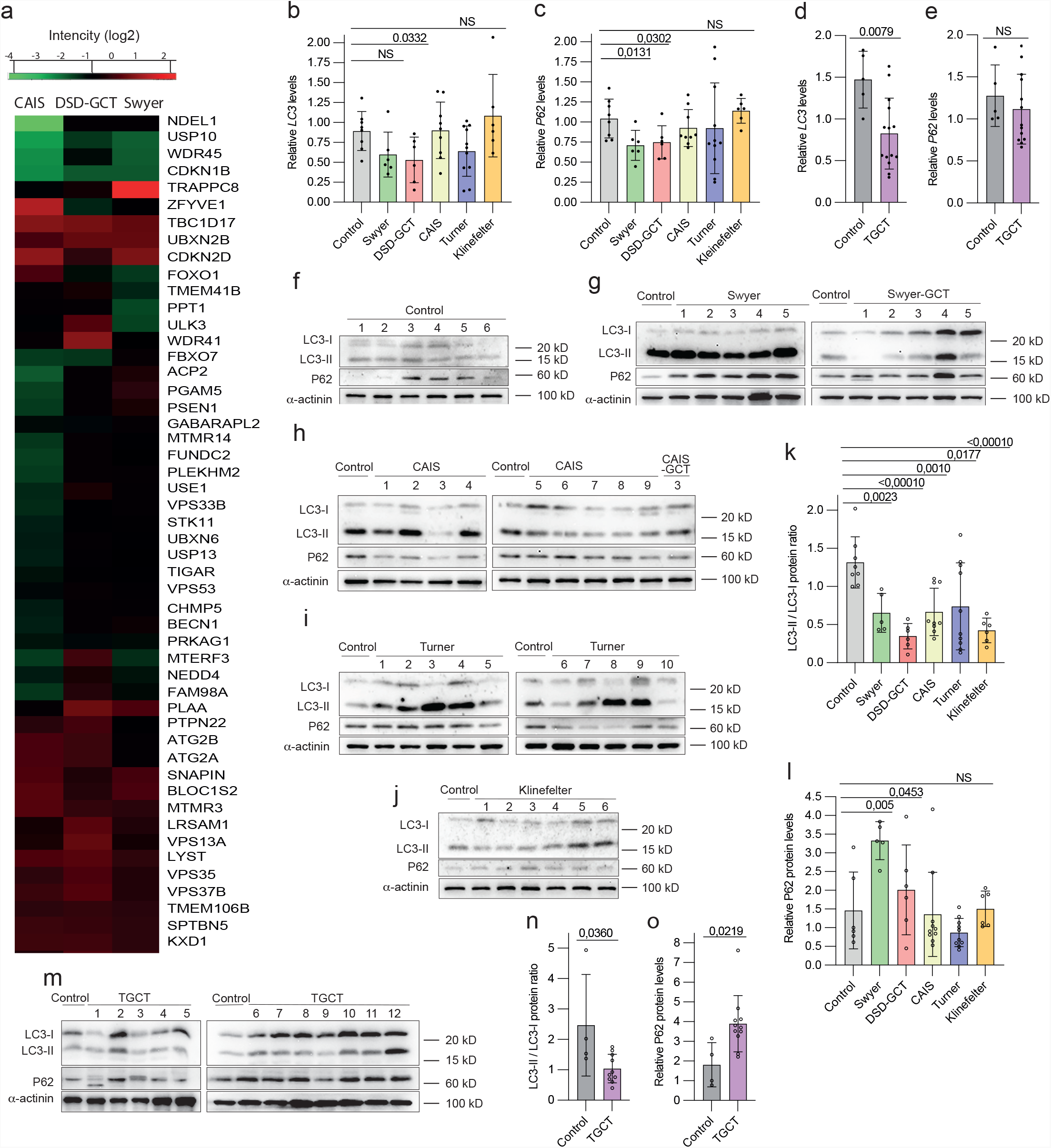
Autophagy deregulation in DSD and TGCT-leukocytes samples. (a) Heat map of GO selection of autophagy specific proteins from entire proteome mass spectrometry data. qRT-PCR analysis of autophagy markers *LC3* and *P63* in patients with DSD (b, c) and TGCT (d, e). Immunoblotting for LC3 and P62 proteins for controls (f), Swyer and Swyer-GCT (g), CAIS and CAIS-GCT (h), Turner (i), Klinefelter (j). Quantification of the ratio of LC3-II to LC3-I (k) and P62 relative protein levels normalized to α-actinin (l) in DSD-patients. Immunoblotting of autophagy markers for TGCT group (m) and quantification (n, o). Unpaired t test was used for statistical analysis, p values are indicated.

We analyzed protein content via measuring ratio of active lipidated LC3-II vs. non-lipidated LC3-I protein forms. We observed in general for DSD-patients a downregulation of LC3-II / LC3-I ratio (Fig. 4f-k). Another autophagy marker, ubiquitin-ligase P62, is normally accumulated, when its protein targets are unable to undergo autophagic degradation (*40*). Accordingly, DSD-samples didn’t show signs of enhanced P62 protein degradation, but rather an accumulation in DSD-GCT and Swyer groups over two-folds (Fig. 4f-j, l). TGCT-samples confirmed these observations, illustrating inhibited LC3-II / CL3-I ratio and accumulation of P62 protein (Fig. 4 m-o). This indicates an inhibition of autophagy flux and consequent accumulation of proteins in leukocytes of DSD and TGCT patients. Therefore, we show that deregulated autophagy and type I interferon signaling supplement DNA damage phenotype in DSD-patients.

### DNA repair mechanisms are altered in leukocytes of DSD-individuals

Deficiency in DNA repair is a leading cause of genetic instability. To understand whether it is relevant to pathology in DSD-patients we analyzed the expression of TP53, a key regulator of DNA damage control(*41*). We observed altered expressions of two TP53 protein variants: full-length (TP53) and its shorter isoform (deltaTP53). We illustrated increase in deltaTP53 expression in all DSD-samples when compared with TP53, except for leukocytes from Turner-patients that also showed statistically significant TP53 upregulation (Fig. 5a-f). The DSD-GCT group failed to upregulate deltaTP53 by four times as it was observed in Swyer, rather resembling relatively small changes in CAIS group in a context of absent DNA damage. Interestingly, TP53 was decreased in leukocytes of DSD-GCT and CAIS-patients (Fig. S3a). DeltaTP53 positively correlates with increasing DNA damage in Swyer and Turner samples defined by γH2AX (Fig. S3b, c). In turn, low deltaTP53 protein levels in DSD-GCT-samples did not correlate with increasing DNA damage (Fig. S3b). Similarly, leukocytes of TGCT-patients showed six-folds increase in deltaTP53, while it failed to correlate with γH2AX (Fig. 5g, h, Fig. S3d). It illustrates that in some TGCT-cases with high DNA damage deltaTP53 fails to be activated similarly to DSD-GCT samples. Therefore, elevated expression of deltaTP53 is intrinsic to samples with genomic instability, while inability to promote deltaTP53 might increase susceptibility to GCT in DSD-patients. Similarly, the entire proteome involved in TP53-regulated DNA repair was generally inhibited (Fig. 5i; Dataset S6). We also observed under expression of particular proteins in DSD-GCT e.g. INF4, USP10, CNOT2, EP300, MIF, PP2R5C, RGCC, EEF1E1.

**Figure 5.**
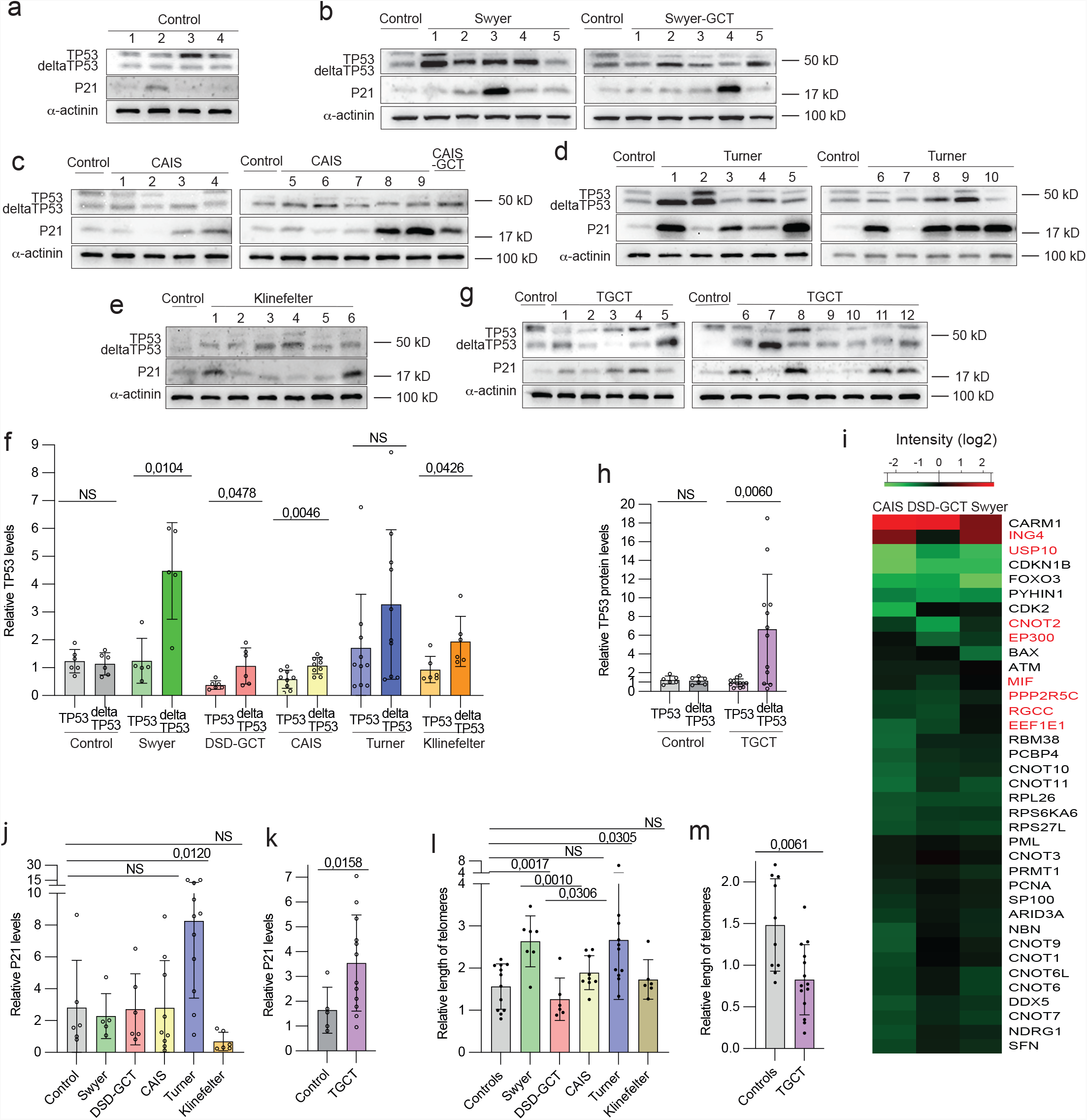
Expression of TP53 protein in leukocytes of DSD- and TGCT-patients. Immunoblotting for TP53 and P21 proteins for controls (a), Swyer and Swyer-GCT (b), CAIS and CAIS-GCT (c), Turner (d) and Klinefelter (e) individuals. (f) Relative quantification full-length (TP53) and depleted isoform (deltaTP53) of TP53 in DSD-groups. Immunoblotting (g) and relative quantification normalized to α-actinin (h) for TGCT-group. (i) Heat map of GO selection of proteins involved in P53-dependent DDR from entire proteome of mass spectrometry analysis. Relative quantification of P21 protein in DSD (j) and TGCT (k) groups. Quantification of relative length of telomeres in DSD (l) and TGCT (m) samples.

To better understand consequences of TP53 protein expression alteration, we examined P21, a TP53 target gene and a key regulator of check-points. We did not detect steady upregulation of P21 among DSD-samples with increased deltaTP53, except for Turner (Fig. 5 a-e, j). However, in TGCT-patients P21 was high, reflecting accumulation of deltaTP53 (Fig. 5g, k). Discrepancy in behavior of deltaTP53 and P21 may indicate a presence of alternative DDR mechanisms in Swyer patients, restricting check-points stimulation upon DNA damage. To additionally characterize DDR mechanisms, we analyzed the length of telomere repeats in different patients’ groups. Deteriorating telomeres can be triggered by aberrant DNA damage repair contributing to genomic instability (*42*). We observed doubled telomere length in Swyer and Turner, implying presence of efficient DNA damage repair mechanisms, which were compromised in DSD-GCT group exposing shortened telomeres (Fig. 5l). Consistently Klinefelter and CAIS groups, with low deltaTP53, did not show an increase in telomeres length. Similar inhibition in telomere length was confirmed in TGCT leukocytes, reflecting distinct status of deltaTP53 in this patient group (Fig. 5m). Taken together we showed deregulated DNA repair mechanisms in leukocytes of DSD samples. Therefore, compromised support form TP53 may underlie their propensity for GCT development.

### TP53 transactivation domain is mutated in DSD-patients with GCT

GCT do not typically carry mutations in TP53 (*43, 44*). To understand the reason behind the inability of DSD-individuals with GCT to induce TP53, we decided to investigate two transactivation domains (TAD) located on the N-terminus. First TAD is encoded on exons 2 and 3 that typically present in full-length TP53. However, when N-terminus is deleted, deltaTP53 only carries second TAD encoded on exon 4. TAD caries crucial Serine (S)/Threonine (T) residues that when phosphorylated block MDM2-dependent degradation of TP53 and promote its binding with other transcriptional factors (*45*). The primary phosphorylation events on S6 and S15 permit consequent post-translational modifications of TP53. In the absence of this phosphorylation TP53 is prone to degradation. Sequencing of the N-terminus of TP53 gene showed missense mutations in exon 2 of Swyer-patients resulting in loss of S/T residues. Only two out of seven studied patients carried both S6 and S15 like in control (Fig. 6a) while exon 4 was mainly unaffected (Fig. 6b), supporting an observation on stabilized deltaTP53 in Swyer-patients (Fig. 5b, f). In turn Swyer-GCT samples had significantly altered sequences of both exons 2 and 4, which could be the reason for decreased stability and so expression of TP53 and deltaTP53 proteins, compared to Swyer without GCT (Fig. 6c, d, Fig. 5b,f). C-terminus modifications can also influence TP53 protein stability(*46*), while we detected no variability in sequences of exons 10 and 11 in Swyer samples, even with GCT (Fig. S4). Therefore, we added an analysis of exons 2 and 4 in other patients’ groups. TAD sequences in exons 2 and 4 mainly remained unaltered in CAIS-samples (Fig. 6e,f). The only exceptions with less S/T in exon 4 were samples CAIS-GCT and CAIS-8, showing hyperplastic Leydig cells by histological analysis (Fig. 6f, Table S1). Turner patients, characterized by low GCT risk, as a rule preserved critical S6 or S15 in exon 2 (Figu. 6g), with couple exceptions that contain S/T on exon 4 and, therefore, are able to stabilize the protein (Fig. 6h). Klinefelter-samples had most of the sequences similar to control (Fig. 6i). Therefore, Swyer-patients with high risk of GCT show evidence of altered TAD sequences in TP53 which correlated with low deltaTP53 expression. While samples of CAIS, Turner and Klinefelter patients without high GCT risk had sequences close to control and, even if TAD on exon 2 altered, it’s usually backed up by unchanged TAD on exon 4 (Fig. 6j).

**Figure 6.**
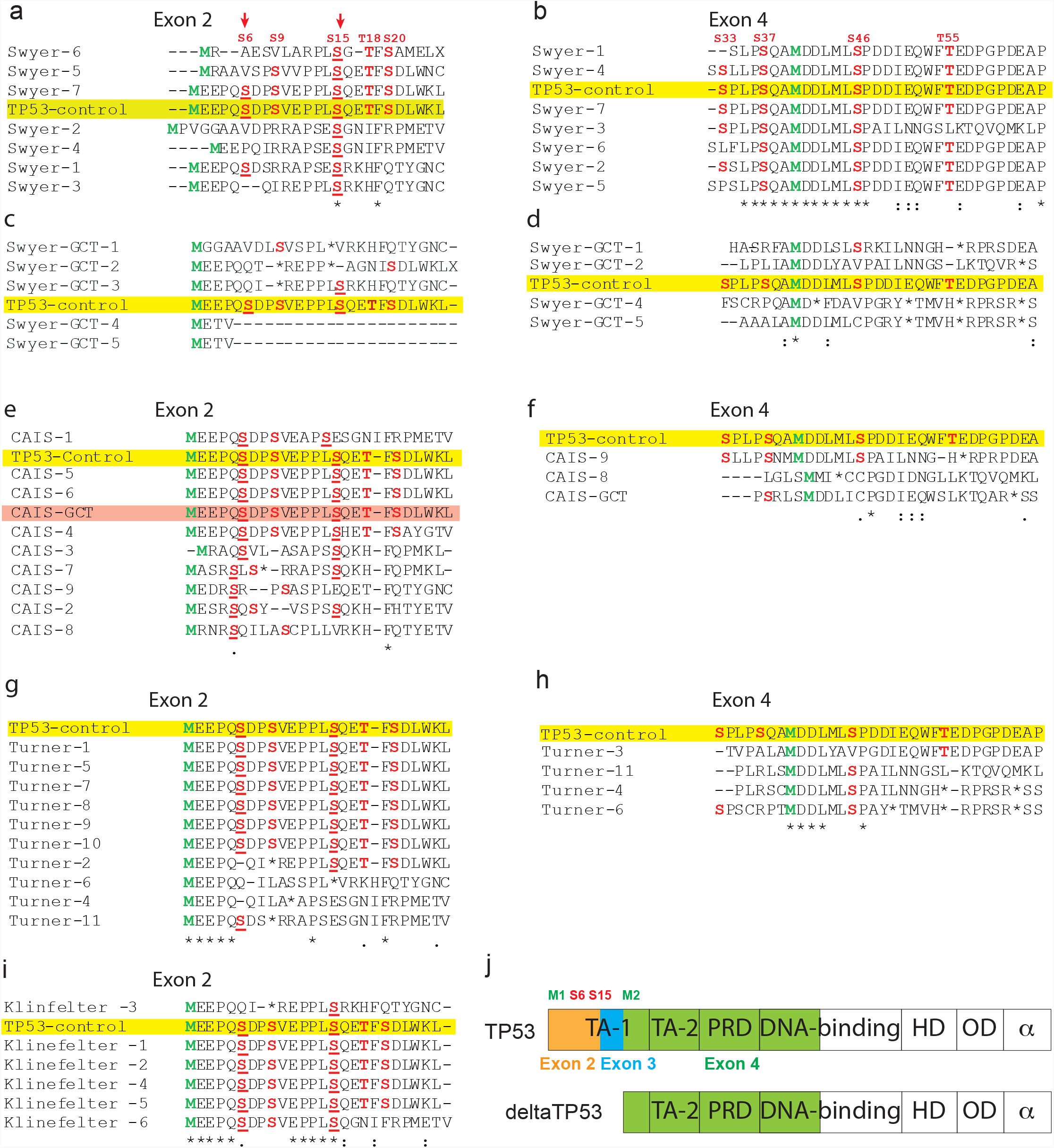
Sequencing of the N-terminus of TP53 gene. Protein sequencing corresponding to transactivation domains were generated based on the DNA sequencing of exons 2 and 4. Protein sequencing of the regions encoded on exon 2 and 4 of Swyer (a, b), Swyer-GCT (c, d), CAIS and CAIS-GCT (e, f), Turner (g, h) and Klinefelter (i) groups. Yellow labels control sample. (j) Scheme illustrating depleted isoform deltaTP53 and a full-length TP53 protein.

### Rescue of DNA damage in DSD-patients

We applied specific drugs to a primary culture of patients’ blood in order to diminish DNA damage. Incubation was for one hour to avoid undesired influence of artificial conditions. First, in order to identify whether decreased autophagy is beneficial for DSD-phenotypes, we supplemented culture medium with Bafilomycin A1, an inhibitor of lysosomal acidity that blocks autophagy at proteolysis stage. As expected we observed low accumulation of LC3 in DSD-samples, as an evidence of decreased autophagy flux compare to control (Fig. 7a, b). γH2AX levels were downregulated upon Bafilomycin A1 addition, implying a necessity of low autophagy rates for DNA repair (Fig. 7a, c). Second, we used enoxacin, known to stimulate DDR and DNA repair via TP53, also in a context of dysfunctional telomeres(*47*), which successfully diminished DNA damage in DSD-leukocytes by two-folds (Fig. 7a,c). Third, we added Idasanutlin, an inhibitor of TP53 and MDM2 interaction, leading to full-length TP53 stabilization (Fig. 7a, d). However, Idasanutlin failed to rescue DNA damage (Fig. 7a,c). This supports our previous observations on the leading role of deltaTP53 in DNA repair mechanisms in DSD-patients. To summarize, we showed that lowering DNA damage levels in DSD-patients’ blood can be done by via autophagy and direct influence on DDR (Fig. 7e).

**Figure 7.**
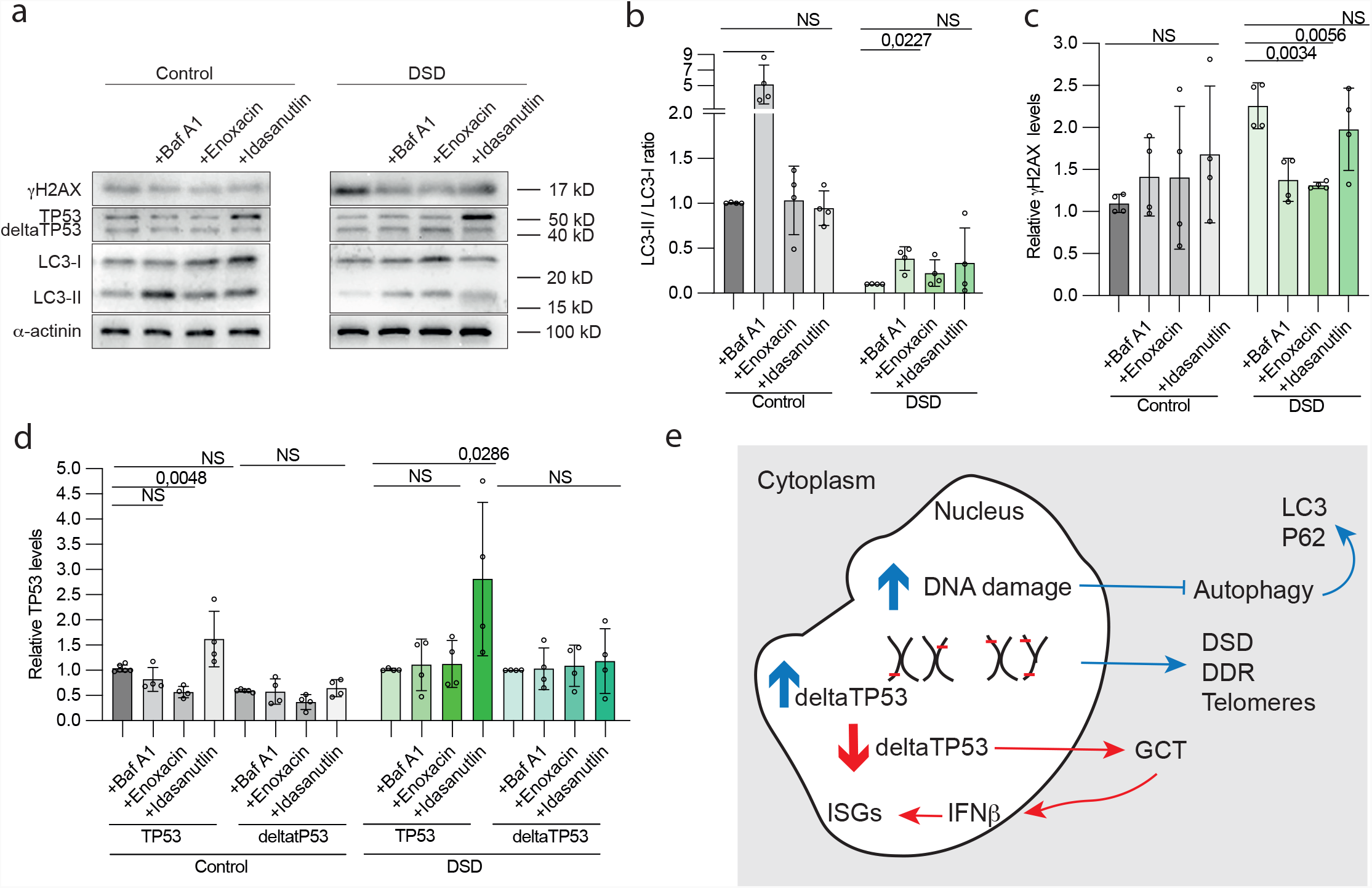
Effects on DNA damage and DDR after short-term culture with drugs. (a) Immunoblotting for TP53, γH2AX and LC3 of control and DSD-samples cultured with Bafilomycin A1 (Baf A1), Enoxacin, Idasanutlin. α-actinin was used as a loading control. (b) Quantification of the ratio of LC3-II vs LC3-I of the immunoblotting (a). Relative protein levels of γH2AX (c) and TP53 (d). (e) A summary scheme illustrating phenotypes specific for DSD (blue arrows) and GCT (red arrows).

## Discussion

In our work we described an increase of DNA damage in leukocytes and gonads of DSD-patients. We also associated DNA damage upregulation with malignancy in dysgenic gonads in different patients’ groups including men with TGCT by evaluating innate immune response activation. Compromised genomic DNA quality suggested an involvement of DNA repair pathways, which we illustrated by upregulation of deltaTP53 in DSD-samples with increased DNA damage. Accordingly, telomere length was upregulated in parallel with increased deltaTP53 expression, illustrating active DNA repair mechanisms in Swyer and Turner patients, while both were decreased in GCT-group. TP53 exhibited a significant increase in missense mutations in a region, encoding transactivation domains and, therefore, compromises protein stability. We also illustrated that leukocytes of DSD-patients are responsive to chemicals regulators, diminishing DNA damage upon direct DNA repair mechanisms activation or autophagy inhibition.

Given that genetic causes for DSD remain unknown for about half of the patients, it’s likely that polygenic mutations or even noncoding regions of genome could be involved(*48*). This supports our idea that a state of genome integrity is behind DSD-phenotypes. DSD-patients often show signs of unstable genome, such as genetic mosaicism with highly variable phenotypes, caused by differences in Y-chromosome breakpoints, unstable idic(Y) chromosomes or dynamic distribution of mosaicism through the organism(*49*). Accordingly, we described Swyer and Turner patients carrying high DNA damage and altered TP53 protein expression that could explain functional or total loss of Y chromosome. Genomic instability in blood of these patients seems to be associated with gonadal dysgenesis. Accordingly, CAIS, caused by mutated androgen receptor, does not have signs of DNA damage, and sometimes is associated with residual germ cells in relatively well-developed gonads. In turn, DNA damage in Klinefelter-patients is likely due to the extra chromosome presence(*27*). Phenotypically males with Klinefelter syndrome possess compromised fertility, however in some cases gonads have residual spermatogenesis, illustrating presence of functional Y chromosomes.

The critical role of genome integrity has been already discussed for germ cells and embryogenesis in humans(*50*). Our data imply a possibility that genotoxic stress in Swyer- or Turner-patients may already be present at the moment of sex tissue determination and, therefore, interfere with gonadal formation due to the defective gene expression from unstable sex chromosomes. Indeed, we observed signs of genome deterioration in misshaped nuclei of cells of so-called streak gonads of Swyer and Turner individuals, while this phenotype was less pronounced in CAIS testicle-like gonads. Genomic instability has been already described in GCT cells of CAIS-patients, however it was associated with mobility of transposable elements (*51*). Considering our findings on increased load of DSBs in blood of patients with DSD, we propose that the state of deterioration of genome integrity likely goes above the transposons activity and rather is a consequence of general misfunction of DNA repair.

TP53 with its high genetic polymorphism is the most commonly mutated gene in cancers (*52*). Full-length protein is known to regulate DDR and being stimulated via N-terminus TAD (*53*). We described a repetitive loss of critical serine residues on N-terminus of TP53 and demonstrated a decrease of TP53 protein in DSD-patients. In turn, deltaTP53 is associated with high DNA damage in Swyer and Turner patients without GCT. Failing to upregulate deltaTP53 seems to increase susceptibility to GCT development in DSD-group. It was shown that suppressed TP53 function causes insensitivity to DDR and continuous cell proliferation, leading to severe shortening of telomeres that normally happens during senescence(*54, 55*). This is likely the reason for decreased telomere length in leukocytes from DSD-GCT group when compared with Swyer and CAIS-patients without GCT. The situation in TGCT was even more evident with shortened telomeres below the controls’ level. Shortened uncapped telomeres tend to fuse, which is accompanied by a whole bunch of genomic instability phenotypes: increased DSBs, chromosomal aneuploidy, micronuclei, misshaped nucleus, chromatin bridges (*56*). In our work we described some of these signs of genotoxic stress in leukocytes and dysgenic gonads, which correlates with TP53 mutation phenotypes. Turner and Swyer both have increased deltaTP53, however only Turner showed P21 induction. This is likely caused by upregulated TP53 in Turner and indicates different situation in Swyer group, possibly relying on mechanisms working against check-points activation. Accordingly, we observed increased length of telomeres in Swyer- and Turner-samples. Discrepancies between TGCT and DSD samples in deltaTP53 regulation show that TGCT samples likely have variable reasons for tumor development. Steady P21 upregulation in TGCT samples also indicates on functional TP53 signaling.

The cell death mechanisms in genotoxic crisis were uncovered only recently and evidently autophagy plays a key role (*40*). We showed for the first time that autophagy and type I Interferon response reflect the state of genomic DNA in DSD- and TGCT-patients’ samples. Accordingly, in leukocytes of patients of studied groups with increased DNA damage, we observed downregulation of autophagy specific gene expression. It is possible that low autophagy prevents apoptosis of cells experiencing genotoxic stress. Indeed, cells with compromised TP53 function and unprotected telomeres are targeted by autophagy (*57*). And cells with knockout of TP53 and ATG7, a key autophagy protein, exhibit intensive proliferation and improved survival rates, since they can bypass senescence. We observed decrease in DNA damage levels in leukocytes from DSD-patients, when autophagy was chemically blocked. It could possibly be connected to the previously described negative influence of autophagy on TP53, and so DNA repair, via proteasomal degradation (*58*). It might be that artificial blockade of autophagy sustained the pull of crucial DDR players for DSD-patients and as a consequence positively influenced on genome status. On the other hand TP53 promotes autophagy specific gene expression(*59*), which could also explain decrease of autophagy in leukocytes of DSD-patients susceptible to TP53 missense mutagenesis. Our data are also in agreement with recently proposed immortalization models, where authors propose loss of TP53 and autophagy function for cells escaping telomere shortening crisis (*57*).

In addition to autophagy, innate immune response is known to be activated by an unstable genome. We observed characteristic for DSD-patients leukocytes upregulation of immune response related proteome with specific changes in samples from patients with GCT. IFNβ signals are stimulated due to the leakage of genomic DNA into the cytoplasm (*27*). Indeed, we detected increased presence of STING protein the key player of cGAS-STING pathways transmitting signals from cytoplasmic dsDNA to the nucleus for type I interferon response stimulation. STING also co-localized with dsDNA in cytoplasm in GCT-cells connecting phenotypes observed in peripheral blood to the actual place of tumorigenesis. In turn, upregulated expression of RNA helicases in DSD-GCT leukocytes samples implies a possibility of RNA induced interferon signaling via similar STING-dependent mechanism (*60*). These finding additionally emphasize an importance to study genome stability in DSD and TGCT patients with high propensity to GCT development. Interestingly, we also observed upregulated *IFNβ* in leukocytes of CAIS-patients, which is likely due to suppressed androgen function (*61*). The role of androgen signaling in GCT regulation and how it is relevant to our patients’ groups is another exciting direction of investigation.

In conclusion we described increased DNA damage phenotype in individuals with differences of sex development and men with testicular germ cell tumors. We defined DDR phenotypes including type I Interferon response and autophagy. DDR regulator TP53 was mutated in samples from patients developing GCT which suggests a new focus for cancer diagnostics and prophylactic treatments to facilitate DSD and GCT-relevant phenotypes.

## Material and methods

### Patients material

Patients with Differences of Sex Development were enrolled into study during routine praxis in the Department of Gynecology Endocrinology and Infertility Disorders. The samples from patients with testicular germ cell tumors were provided us by Urology Department. The samples were collected with written patients consent. For this study we analyzed 40 patients with DSD and 19 samples with GCT. DSD-patients included individuals with Swyer, Turner, Klinefelter and Complete Androgen Insensitivity (CAIS) Syndromes. The patients are listed with additional information (Table S1 and S2). Patient sample undergo standard characterization procedure in genetics and histological facility of Heidelberg University Hospital.

### Leukocytes isolation from the peripheral blood

Blood was collected in EDTA and subjected to leukocytes isolation according to the stablished protocol. 10 ml of blood was mixed with 30 ml of lysis buffer (155 mM NH_4_Cl, 10 mM KHCO3, 0,1 mM EDTA ph 7.4) and incubated 30’ on ice. Then mix was centrifuged 10’ at 1200 rpm at 4°C. The pellet was washed three times with 10 ml lysis buffer and then subjected for DNA, RNA and protein isolation with NucleoSpin® TriPrep kit (Marcherey-Nagel).

### Western blotting

Protein from leukocytes was analyzed via Western Blot analysis following modified protocol(*62*). The protein concentration was determined by Protein Quantification Assay (Cat.# 740967,Macherey-Nagel). 10 µg of the protein was separated by SDS-PAGE electrophoresis, then transferred onto PVDF membrane (Immobilon-P membrane (Cat.# IPVH00010, Millipore). The membrane was blocked with 5% skim milk (Cat. # T145, Carl Roth) or 3 % BSA (Cat. # 8076.4, Carl Roth) 1 h at RT. Then incubated at 4°C overnight with primary antibodies. After washing three times with buffer TNT (50 mM Tris, 150 mM NaCl, 5 mM EDTA and 0.05% Tween 20; pH 7.6), the membrane was incubated with corresponding secondary antibodies (peroxidase conjugated Goat Anti-Rabbit IgG (Cat.# 111-035-046, Dianova) or peroxidase conjugated Goat Anti-Mouse IgG (Cat.# 115-035-062, Dianova) for 1 h at RT. Protein bands were visualized by enhanced chemiluminescence with an ECL kit (SuperSignal West Femto Substrat (Cat. # 34095, Thermo Scientific). Image development was performed on iBright FL1000 Imager (Cat.# A32748, Invitrogen). Antibodies list is provided (Table S3). Every point in the plots of data for WB analysis represents one individual patient sample or experimental measurement, total sample size is represented by 3-10 points on the graph.

### Quantitative real-time PCR

1 µg RNA was used for cDNA synthesis with Superscript II (Cat.# 18064022, Invitrogen). qPCR analysis was performed on an ABI Prism 7900HT Fast Real-Time PCR System (Applied Biosystems). The cycling conditions were as follows: 50°C for 2 min, denaturation at 95°C for 10 min followed by 40 cycles at 95°C for 15 s, and a combined annealing and extension step at 60°C for 60 s. The list of TaqMan assays (Termofisher) is provided (Table S4). Every point in the plots of data for qRT-PCR analysis represents one individual patient sample or experimental measurement, total sample size is represented by 3-10 points on the graph.

### Blood culture with drugs

For each condition we prepared 10 cm dishes with 10 ml culture medium, containing 2 ml of blood, collected in citrate and containing approximately 5×10^6 leukocytes per 1 ml; 15% FCS (Cat. #10500064, Life Technologies); 20 µl PHA-L (Cat.# 00-4977-03, Life Technologies). We added Enoxacin (10µM, S1756, Selleckchem), Idasanutlin (10µM, HY-15676, Hycultec), Bafilomycin A1 (100nM, 11038, Cayman). DMSO was used as the control. After 1 h incubation at 37°C, 5%CO2 cells were collected, washed 2 times with 1xPBS (Cat. # 14190-094, Gibco) and subjected to leukocytes isolation procedure as described above. DNA, RNA and protein isolation from the leukocytes was performed with NucleoSpin® TriPrep kit (Marcherey-Nagel).

### Hematoxylin / eosin staining and microscopy

Tissue biopsies were fixed in buffered formaldehyde and subsequently embedded in paraffin. Sections, 4 µm thick, were stained with the standard protocol for Hematoxylin / Eosin. Immunohistochemistry for AMH was performed as previously published (*12*). The imaging was performed using light microscope (LEICA) with magnification 100x. The image analysis was performed using ImageJ software. The circularity and roundness parameters were plotted for statistical analysis using GraphPad.

### Immunofluorescence

Immunostaining was performed on 4 μm dehydrated FFPE tissue sections. Antigen retrieval was performed by immersing the slides in 10 mM citrate buffer (pH 6.0) in a steamer for 10 min. Next, tissue sections were allowed to cool and then kept at room temperature for 20 min. Slides were washed twice in PBS and three times in washing buffer (PBS containing 0,1 % Triton X-100). Tissue sections were incubated in 3% H_2_O_2_/Methanol for 10 min. Sections were washed three times with washing buffer and then blocked with blocking buffer (5% goat serum (cat. # S-1000, Vector) in washing buffer for 1 h at room temperature. Tissue sections were incubated with primary antibody in a humid box at 4 °C for 16 h. The primary was diluted in 1% goat serum/washing buffer. After three washes in washing buffer secondary fluorescence-conjugated antibodies (Goat Anti-Rabbit/Mouse DyLight 488 conjugated, cat # 35552, Thermo Scientific) was diluted to a concentration of 2 µg/ml in 1% goat serum/washing buffer and incubated for 2 h at room temperature in a dark humid box. Slides were then rinsed in washing buffer five times for 3 min each and nuclei were stained with 4lr,6-diamidino-2-phenylindole (DAPI, cat # D-1388, Sigma) at a concentration of 1 μg/ml for 10 min at RT. Sections were then rinsed three times with water for 3 min each before being mounted and coverslips were applied using ProLong Glass Antifade Mountant (cat. #P36980, Invitrogen). The imaging was performed using a fluorescence microscope (LEICA, DMLB), using a HQ camera SpectraCube (Applied Spectral Imaging, Mannheim, Germany) and a 20x-air, 63x-oil or 100x-oil objective (PL FLUOTAR 20x/0,5 PH2, HCX PL APO 63x/1,32-0,6 OIL, PL FLUOTAR 100x/1,3 OIL PH3) under the control of the Spectral imaging acquisition software 2.6 (Applied Spectral Imaging, Germany). For immunofluorescent analysis, 100-1000s of cells were analyzed using automatic software ImageJ. The means of each measurement/experiment were calculated and plotted for presentation and for statistical analysis using GraphPad.

Primary antibodies used anti-γ-H2AX rabbit polyclonal antibody (0,25μg/mL, cat.#ab11174, Abcam), anti-STING (D2P2F) rabbit monoclonal (1/50 dilution, 13647S, Cell Signaling Technology), anti-dsDNA mouse monoclonal (1/50 dilution, 58749, Santa Cruz Biotechnology), anti-OCT4 mouse monoclonal (1/300 dilution, cat.#SC-5279, Santa Cruz Biotechnology).

Gonadal tissue sections were provided by the tissue bank of the National Centre for Tumor Diseases (NCT, Heidelberg, Germany) in accordance with the regulations of the tissue bank and the approval of the ethics committee of Heidelberg University.

### Telomere length assay

Genomic DNA was isolated from blood according the previously established protocol. In short, 10 ml of blood collected in EDTA was mixed with 30 ml of lysis buffer (155 mM NH_4_Cl, 10 mM KHCO3, 0,1 mM EDTA pH7.4), incubated on ice for 30’ and centrifuged for 10’ at 1200 rpm, 4°C. The pellet was washed three times with 10 ml of lysis buffer and resuspended in 5 ml of SE-Buffer (75 mM NaCl, 25 mM EDTA pH 8.0), supplemented with 250 µl of 20% SDS and 40 µl Proteinase K (10 mg/ml) and left overnight at 37°C. Salt precipitation was done by repetitive mix with 15 ml with SE-Buffer, supplemented with 3,3 ml of saturated NaCl and centrifugation for 15’ at RT until complete removal of the pellet. Next, DNA precipitation was performed with 1,7 ml 3 M Sodium-Acetat pH7.0 and 17 ml Isopropanol. After DNA-fiber could be collected, wash them with 80% Ethanol. Let the fiber dry and dissolve them in TE (10 mM Tris, 1 mM EDTA pH 8.0). We used 5µg of genomic DNA per reaction for Relative Human Telomere Length Quantification qPCR Assay Kit (Cat. #8908, ScienCell). Every point in the plots of data for qRT-PCR analysis represents one individual patient sample, total sample size is represented by 3-10 points on the graph.

### Sequencing

We used DSD-patient’s genomic DNA for sequencing. We analyzed exons 2-3 and exon 4 of P53 gene according to the established protocol (IARC, 2019). The primers used for exons 2-3 F-tctcatgctggatccccact; R-agtcagaggaccaggtcctc; exon 4 F-tgctcttttcacccatctac; R-atacggccaggcattgaagt. The sequencing was performed using BigDye Terminator v1.1 Cycle Sequencing Kit (Applied Biosystems, Cat. no. 4337450) and the ABI PRISM 3100 Genetic Analyzer. Sequencing Data that support the findings of this study have been deposited into open access NCBI GeneBank with the accession codes for ON398073 - ON398110 (TP53-exon2) and ON398111 - ON398128 (TP53-exon4).

### Mass spectrometry

Proteome-wide expression profiling of leukocytes from three Swyer, CAIS, DSD-GCT patients and fertile controls, was supported by the Core Facility for Mass Spectrometry & Proteomics (CFMP) at the Center for Molecular Biology at University Heidelberg (ZMBH). 50µg of protein was precipitated using Wessel-Flügge method(*63*) and protein pellet was dissolved in 8M Urea buffer. After reduction and alkylation for 30 minutes at RT using 10 mM TCEP and 40 mM CAA, Lys-C was added at 1:100 enzyme:protein ratio and incubated for 4h at 37° C. Mixture was diluted 1:4 with 50 mM TEAB pH 8.5 and trypsin was added in ratio 1:50. After overnight incubation at 37° C, samples were acidified, desalted on self-made C18 Empore® extraction discs (3M) StageTips(*64*), concentrated in a SpeedVac and stored at −20° C until measured.

Samples were suspended in 0.1% TFA and an equivalent to 1 µg of peptides was analyzed using Ultimate 3000 liquid chromatography system coupled to an Orbitrap QE HF (Thermo Fisher). An in-house packed analytical column (75 µm x 200 mm, 1.9 µm ReprosilPur-AQ 120 C18 material Dr. Maisch, Germany) was used. Mobile phase solutions were prepared as follows, solvent A: 0.1% formic acid / 1% acetonitrile, solvent B: 0.1% formic acid, 89.9% acetonitrile.

Peptides were separated in a 120 min linear gradient started from 3% B and increased to 23% B over 100 min and to 38% B over 20 min, followed by washout with 95% B. The mass spectrometer was operated in data-dependent acquisition mode, automatically switching between MS and MS2. MS spectra (m/z 400–1600) were acquired in the Orbitrap at 60,000 (m/z 400) resolution and MS2 spectra were generated for up to 15 precursors with normalized collision energy of 27 and isolation width of 1.4 m/z.

The MS/MS spectra were searched against the Swiss-Prot Homo sapiens protein database (UP000005640, June 2020, 20531 sequences) and a customized contaminant database (part of MaxQuant, MPI Martinsried) using Proteome Discoverer 2.5 with Sequest HT (Thermo Fisher Scientific). A fragment ion mass tolerance was set to 0.02 Da and a parent ion mass tolerance to 10 ppm. Trypsin was specified as enzyme. Carbamidomethyl was set as fixed modification of cysteine and oxidation (methionine), acetylation (protein N-terminus) and methionine loss (protein N-terminus) as variable modifications. Peptide quantification was done using precursor ion quantifier node with Top N Average method set for protein abundance calculation and N for Top N set to 3.

For comparison of each patient group with the control we calculated the protein levels fold change ratios. The same datasets were used to analyze DDR, autophagy- and immune response, and P53-specific proteome. Presence of at least one unique peptide is required for identification of reported protein groups. Minimum of two ratio counts were required for the quantitation based only on unique and razor peptides. For details, see the Datasets S1-S6.

The proteomics data used in this manuscript are deposited in the ProteomeXchange Consortium (http://proteomecentral.proteomexchange.org) via the PRIDE partner repository and can be accessed with a code PXD033635.

## Acknowledgements

We would like to thank Core Facility for Mass Spectrometry & Proteomics of University of Heidelberg for mass spectrometry analysis. Also, we would like to thank Prof. E. Alexandrova for the critical comments on the manuscript and TP53 work in particular.

## Author contributions

MK initiated and supervised the project, performed experiments, analyzed data, supervised JZ and AS, and wrote the paper. JZ performed the experiments and analyzed the data. AS did telomere assay. PFH, JR, MH, MB, TS provided the patients material. ML performed mass spec analysis. TS obtained the funding. All authors commented on the paper.

