## Supplementary material for "Genomic instability in patients with sex determination defects and germ cell cancer": Suppl. files: Supplementary figure 1.pdf

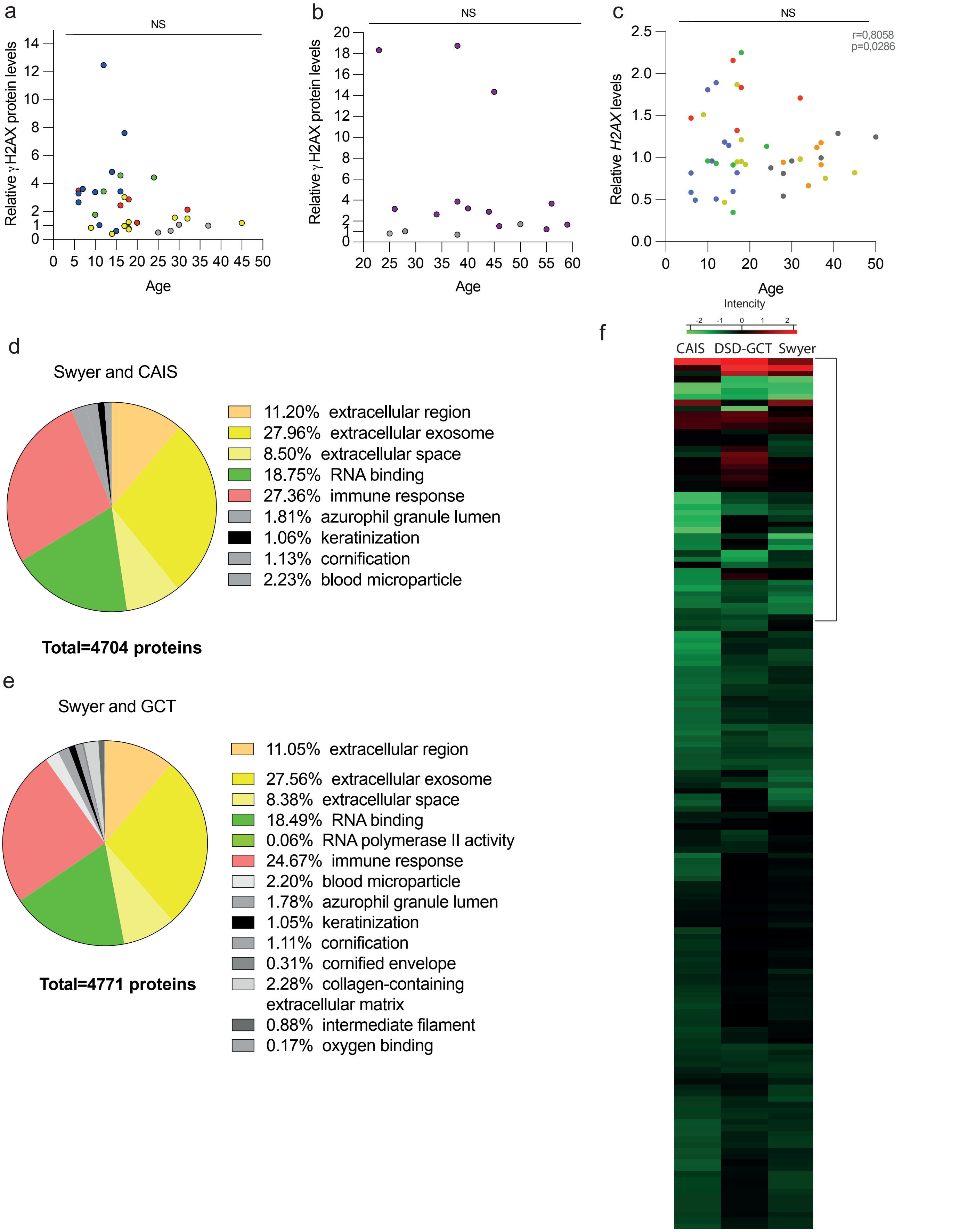


**Supplementary figure 1. Changes in DNA damage markers expression in leukocytes with age in DSD and TGCT-patients.** γH2AX relative protein expression levels from DSD (a) and TGCT (b) samples groups correlated with age. (c) *H2AX* relative transcripts levels were plotted to correlate with age for DSD-patients. Pearson's correlation test was applied for statistical analysis and was positive for control group in (c). Colored dots correspond to different DSD-groups: Swyer-green, DSD-GCT-red, CAIS-yellow, Turner-blue; and control group - grey. Correlation coefficient and p values are indicated. All other cases didn't have statistically significant correlation. The whole proteome mass spectrometry analysis data are represented from three Swyer and three CAIS (d) or DSD-GCT (e) patients. Green labeled group of proteins involved in RNA binding pathways was downregulated, while all others were upregulated, when normalized to the control. (f) GO on DDR selection shows mostly inhibited gene expression. The area indicated was chosen for magnified selection for Figure 1m.
