## Supplementary material for "Genomic instability in patients with sex determination defects and germ cell cancer": Suppl. files: Supplementary figure 2.pdf

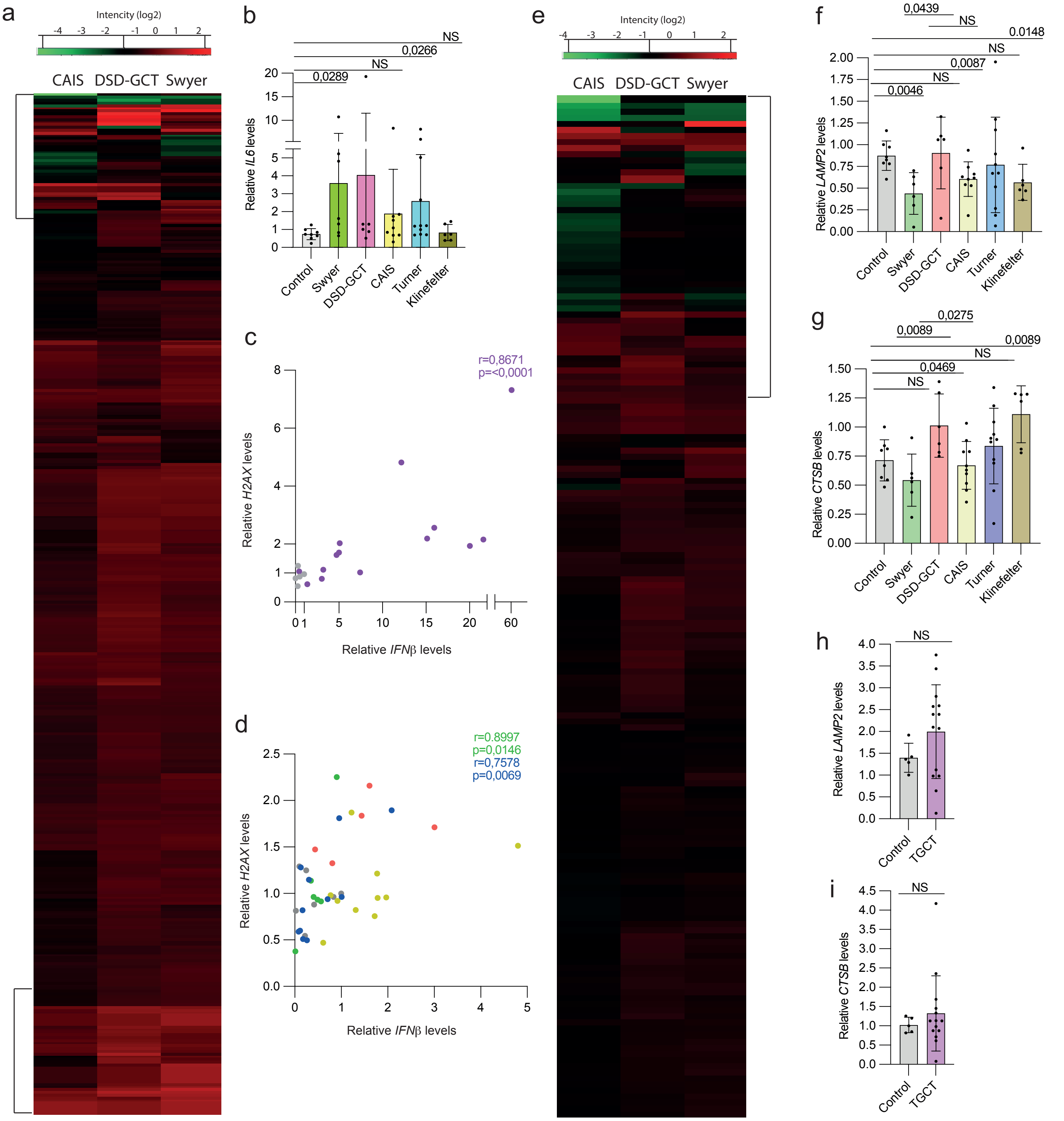


**Supplementary figure 2. DDR in leukocytes of DSD and TGCT-patients.** Heat maps representing selection of proteins involved in immune response (a) and autophagy (e) from mass spectrometry analysis data. Selected area in (a) is magnified in a figure 3a, b, and selection in (e) is magnified in a figure 4a. qRT-PCR data of the NF-kB pathway target gene expression *IL6* in DSD-group (b). Correlation of DNA damage marker *H2AX* and innateimmune specific gene *IFNβ* in TGCT (c) and DSD (d) groups (Swyer-green, DSD-GCT-red, CAIS-yellow, Turner-blue, control-grey). qRT-PCR data of lysosomal gene expression markers *LAMP2* and *CTSB* in DSD (f, g) and TGCT (h, i) samples. Pearson's correlation coefficients and p values after unpaired t-test are indicated.
