## Supplementary material for "Genomic instability in patients with sex determination defects and germ cell cancer": Suppl. files: Supplementary figure 3.pdf

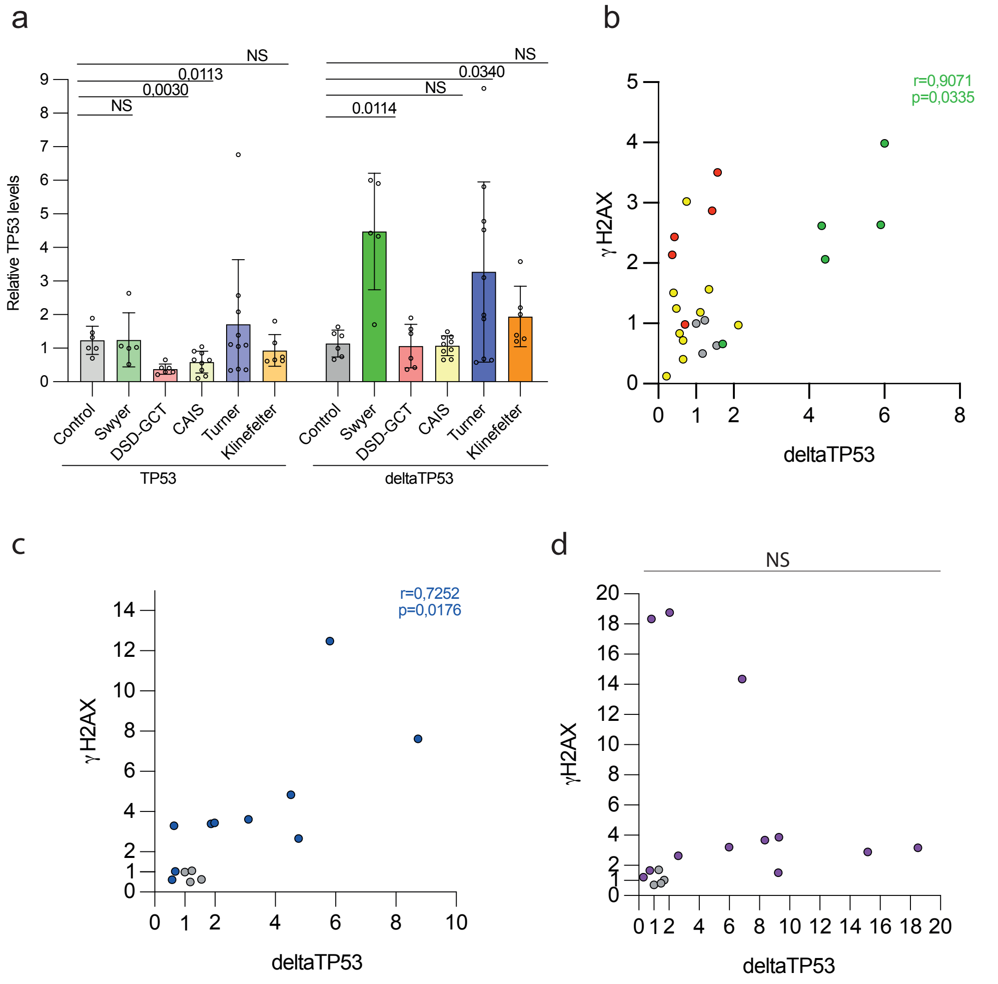


**Supplementary figure 3. Associated TP53 expression with DNA damage levels.** (a) Quantification of TP53 protein levels relative to α-actinin from immunoblotings (Figure 4 a-e). Pearson's correlation was performed for DSD-samples (b, c). CAIS (yellow), DSD-GCT (red) and control (grey) groups didn't show significant correlation (b) as well as TGCT-group (d). Swyer (green) and Turner (blue)-samples showed significant correlation between γH2AX and TP53. Pearsons' correlation coefficients and p values from unpaired t test are indicated.
