## Supplementary material for "Genomic instability in patients with sex determination defects and germ cell cancer": Suppl. files: Supplementary figure 4.pdf

a

Exon 10

```
Swyer-GCT-1    ---CGGCGGGTTATCCTTCTTCCCCCCCCTGTTGCTGCAGATCGTGGGCGTGAGCGCTTC  57
Swyer-GCT-4    --AAGCCGGTATATCCTTCTTCCCCCCCCTGTTGCTGCAGATCGTGGGCGTGAGCGCTTC  58
TP53-control   -----------------------------------GCAGATCCGTGGGCGTGAGCGCTTC  25
Swyer-1        GGCTGGCAGGTTATCTTCGTTCCCCCCCCTGTTGCTGCAGATCGTGGGCGTGAGCGCTTC  60
Swyer-2        --TGCAGCCGGTTTCTTCGTTCCCCCCCCTGTTGCTGCAGATCGTGGGCGTGAGCGCTTC  58
                                                          *****************

Swyer-GCT-1    GAGATGTTCCGAGAGCTGAATGAGGCCTTGGAACTCAAGGATGCCCAGGCTGGGAAGGAG  117
Swyer-GCT-4    GAGATGTTCCGAGAGCTGAATGAGGCCTTGGAACTCAAGGATGCCCAGGCTGGGAAGGAG  118
TP53-control   GAGATGTTCCGAGAGCTGAATGAGGCCTTGGAACTCAAGGATGCCCAGGCTGGGAAGGAG  85
Swyer-1        GAGATGTTCCGAGAGCTGAATGAGGCCTTGGAACTCAAGGATGCCCAGGCTGGGAAGGAG  120
Swyer-2        GAGATGTTCCGAGAGCTGAATGAGGCCTTGGAACTCAAGGATGCCCAGGCTGGGAAGGAG  118
               ************************************************************

Swyer-GCT-1    CCAGGGGGGAGCAGGGCTCACTCCAGGTGAGTGACCTCAGCCCCTTCCTGGCCCTACTCC  177
Swyer-GCT-4    CCAGGGGGGAGCAGGGCTCACTCCAGGTGAGTGACCTCAGCCCCTTCCTGGCCCTACTCC  178
TP53-control   CCAGGGGGGAGCAGGGCTCACTCCAGGT--------------------------------  113
Swyer-1        CCAGGGGGGAGCAGGGCTCACTCCAGGTGAGTGACCTCAGCCCCTTCCTGGCCCTACTCC  180
Swyer-2        CCAGGGGGGAGCAGGGCTCACTCCAGGTGAGTGACCTCAGCCCCTTCCTGGCCCTACTCC  178
               ****************************
```

b

Exon 11

```
TP53-control ----------------------AGCCACCTGAAGTCCAAAAAGGGTCAGTCTACCTCCCG    38
Swyer-GCT-1  ----GCTCTCTCCTCCTTTGCTCCTCAGCCACTGAAGCCAAAAAGGGCAGCTACCTCCCG    56
Swyer-GCT-4  GTCCTCTCCCTCCTCTTTGCTCCTCAGCCTCCTGAAGCCAAAAAGGGCAGCTACCTCCCG    60
                                         *    *     ***  *     **********

TP53-control CCATAAAAAACTCATGTTCAAGACAGAAGGGCCTGACTCAGACTGACATTCTCCACTTCT    98
Swyer-GCT-1  CCATAAAAAACTCATGTTCAAGACAGAAGGGCCTGACTCAGACTGACATTCTCCACTTCT    116
Swyer-GCT-4  CCATAAAAAACTCATGTTCAAGACAGAAGGGCCTGACTCAGACTGACATTCTCCACTTCT    120
             ************************************************************

TP53-control TGTTCCCCACTGACAGCCTCCCACCCCCATCTCTCCCTCCCCTGCCATTTTGGGTTTTGG    158
Swyer-GCT-1  TGTTCCCCACTGACAGCCTCCCACCCCCATCTCTCCCTCCCCTGCCATTTTGGGTTTTGG    176
Swyer-GCT-4  TGTTCCCCACTGACAGCCTCCCACCCCCATCTCTCCCTCCCCTGCCATTTTGGGTTTTGG    180
             ************************************************************

TP53-control GTCTTTGAACCCTTGCTTGCAATAGGTGTGCGTCAGAAGCACCCAGGACTTCCATTTGCT    218
Swyer-GCT-1  GTCTTTGAACGCGTTGCAATGCAATGGGGTGTGCGTCAAA--------------------    216
Swyer-GCT-4  GTCTTTGAACCGCGTTGCATATGCAATGGTGTGCGTCAAA--------------------    220
             **********                 *     *   *
```

**Supplementary figure 4. DNA sequence alignment.** Genomic DNA of two Swyer-GCT samples were sequenced on the regions encoding exon 10 (a) and exon 11 (b) of gene TP53.
