## Supplementary material for "Genomic instability in patients with sex determination defects and germ cell cancer": Suppl. files: Supplementary table 1 DSD.pdf

Supplementary table 1.
List of DSD-patients.

| | Patient groups | Age | Karyotype | Malignant Tumor (gonadal biopsy) | Gonadal tissue histology |
|---|---|---|---|---|---|
| | Swyer-GCT | | | | |
| 1 | 1 | 16 | 46, XY | Dysgerminoma | CD117+, PLAP+ |
| 2 | 2 | 18 | 46, XY | Seminoma/ Dysgerminoma | PAS+ |
| 3 | 3 | 17 | 46, XY (SRY mutation) | Dysgerminoma | CD117+, D2-40+, OCT4+, PLAP+, SALL4+ |
| 4 | 4 | 32 | 46, XY | Dysgerminoma, Gonadoblastoma | AFP+, β-HCG+ |
| 5 | 5 | 6 | 46, XY | Dysgerminoma | KiA10+, AP + |
| | Swyer | | | | |
| 6 | 1 | 24 | 46, XY | Not found | |
| 7 | 2 | 16 | 46, XY | Not found | |
| 8 | 3 | 12 | 46, XY | Not found | |
| 9 | 4 | 10 | 46, XY | Not found | |
| 10 | 5 | 18 | 46, XY | Not found | |
| 11 | 6 | 16 | 46, XY | Not found | |
| 12 | 7 | 17 | 46, XY | Not found | |
| | CAIS | | | | |
| 13 | 1 | 29 | 46, XY | Not found | Negative for AFP, OCT4, SALL4, PLAP, S100, CD117, beta-HCG |
| 14 | 2 | 45 | 46, XY | Not found | Negative for S100, SALL4, AFP, C2-40, OCT4, PLAP, beta-HCG |
| 15 | 3 | 14 | 46, XY | Not found | |
| 16 | 4 | 18 | 46, XY | Not found | |
| 17 | 5 | 9 | 46, XY | Not found | |
| 18 | 6 | 17 | 46, XY | Not found | |
| 19 | 7 | 17 | 46, XY | Not found | |
| 20 | 8 | 18 | 46, XY | Not found | hyperplastic Leydig cells |
| 21 | 9 | 32 | 46, XY | Not found | Differentiated Sertoli-Leydig cells tumor, secondary Sertoli cells hyperplasia, no germ cells tumor, no malignancy<br>OCT4+, TSPY+, DDX3Y+ |
| | CAIS-GCT | | | | |
| 22 | 1 | | 46, XY | Dysgerminoma/ Gonadoblastoma | D2-40, OCT4, SALL4 |
| | Turner | | | | |
| 23 | 1 | 14 | 45, X0 | No biopsy | |
| 24 | 2 | 17 | 45, X0 | No biopsy | |
| 25 | 3 | 10 | 45, X0 | No biopsy | |
| 26 | 4 | 12 | 45, X0 | No biopsy | |
| 27 | 5 | 16 | 45, X0 | No biopsy | |
| 28 | 6 | 15 | 45, X0 | No biopsy | |

| | | | | | |
|---|---|---|---|---|---|
| 29 | 7 | 11 | 45, X0 | No biopsy | |
| 30 | 8 | 7 | 45, X0 | No biopsy | |
| 31 | 9 | 6 | 45, X0 | No biopsy | |
| 32 | 10 | 6 | 45, X0 | No biopsy | |
| 33 | 11 | 12 | 45, X0 | No biopsy | |
| | Klinefelter | | | | |
| 34 | 1 | 34 | XXY | Not found | pronounced fibrosis |
| 35 | 2 | 28 | XXY | Not found | |
| 36 | 3 | 37 | XXY | Not found | |
| 37 | 4 | 32 | XXY | Not found | pronounced fibrosis |
| 38 | 5 | 37 | XXY | Not found | |
| 39 | 6 | 36 | XXY | Not found | reactive Leydig cell hyperplasia |
