## Supplementary material for "Genomic instability in patients with sex determination defects and germ cell cancer": Suppl. files: Supplementary table 2 TGCT.pdf

Supplementary table 2.
List of TGCT-patients.

| TGCT | Age | Karyotype | Tumor |
|---|---|---|---|
| 1 | 38 | 46, XY | Seminoma |
| 2 | 23 | 46, XY | Seminoma;<br>Embryonal carcinoma;<br>Yolk sac tumor;<br>intratubular germ cell neoplasia unclassified |
| 3 | 55 | 46, XY | Seminoma |
| 4 | 59 | 46, XY | Embryonal carcinoma;<br>Teratoma |
| 5 | 34 | 46, XY | Seminoma |
| 6 | 38 | 46, XY | Seminoma, |
| 7 | 26 | 46, XY | Seminoma;<br>Testicular intraepithelial neoplasia |
| 8 | 56 | 46, XY | Seminoma |
| 9 | 45 | 46, XY | Seminoma |
| 10 | 40 | 46, XY | Seminoma |
| 11 | 46 | 46, XY | Seminoma |
| 12 | 44 | 46, XY | Seminoma |
| 13 | 32 | 46, XY | Embryonic Carcinoma<br>Teratoma;<br>Chorionic Cancer;<br>Seminoma |
| 14 | 25 | 46, XY | Seminoma;<br>Embryonal Carcinoma;<br>Yolk sac Tumor;<br>Chorionic Carcinoma |
