## Supplementary material for "Genomic instability in patients with sex determination defects and germ cell cancer": Suppl. files: Supplementary table 3 abs.pdf

List of antibodies.

| Antibody | Concentration/dilution used | Cat. No | Company |
|---|---|---|---|
| γH2AX | 1 mg/ml; 1/5000 | ab11174 | Abcam |
| H2AX | 1,078 mg/ml; 1/5000 | ab124781 | Abcam |
| α-actinin | 200 μg/ml; 1/30.000 | SC-17829 | Santa Cruz Biotechnology |
| dsDNA | 1/50 | SC-58749 | Santa Cruz Biotechnology |
| LC3 | 1/500 | #3868 | Cell Signaling |
| OCT4 | 1/300 | SC-5279 | Santa Cruz Biotechnology |
| P62 | 1/500 | #5114 | Cell Signaling |
| P53 | 1/500 | M7001 | Agilent (DAKO) |
| P21 | 1/500 | #2947 | Cell Signaling |
| STING | 1/50 | 13647S | Cell Signaling |
