## Supplementary material for "Genomic instability in patients with sex determination defects and germ cell cancer": Suppl. files: Supplementary table 4 TaqMan.pdf

List of TaqMan assays.

| Target gene name | Reference |
|---|---|
| H2AFX | Hs00266783_s1 |
| LC3 | Hs00261291_m1 |
| P62 | Hs00177654_m1 |
| IFNβ | Hs01077958_s1 |
| ISG15 | Hs00192713_m1 |
| ISG56 | Hs01675197_m1 |
| IL6 | Hs00174131_m1 |
| LAMP2 | Hs00903582_m1 |
| CTSB | Hs00947439_m1 |
| Endogenous control gene name | |
| TBP | Hs00427620_m1 |
| HPRT | Hs99999909_m1 |
| GAPDH | Hs02758991_g1 |
| RPLPO | Hs99999902_m1 |
